# High-Speed AFM directly visualizes conformational dynamics of the HIV-Vif protein complex

**DOI:** 10.1101/2020.05.18.102749

**Authors:** Yangang Pan, Luda S. Shlyakhtenko, Yuri L. Lyubchenko

## Abstract

Viral infectivity factor (Vif) is a protein that is essential for the replication of the HIV-1 virus. The key function of Vif is to disrupt the antiviral activity of APOBEC3 proteins, which mutate viral nucleic acids. Inside the cell, Vif binds to the host cell proteins Elongin-C, Elongin-B, and CBF-β, forming a four-protein complex called VCBC. The structure of VCBC in complex with the Cullin5 (Cul5) protein has been solved by X-ray crystallography, and recently, using molecular dynamic (MD) simulations, the dynamics of VCBC and VCBC-Cul5 complexes were characterized. Here, we applied time-lapse high-speed atomic force microscopy (HS-AFM) to visualize the conformational changes of the VCBC complex. We determined the three most favorable conformations of the VCBC complex, which we identified as triangle, dumbbell, and globular structures. In addition, we characterized the dynamics of each of these structures. While our data show a very dynamic behavior for all these structures, we found the triangle and dumbbell structures to be the most dynamic. These findings provide insight into the structure and dynamics of the VCBC complex and support further research into the improvement of HIV treatment, as Vif is essential for virus survival in the cell.

## Introduction

HIV is an enveloped retrovirus that causes AIDS in humans (1). The virus particle contains two identical RNA copies and structural and replication enzymes for virus reproduction (2). One of the critical proteins needed for the virus survival is viral infectivity factor (Vif) (3). Vif, a small and unstructured 23 kDa protein (4,5), counteracts the potent HIV-1 inhibitors, human A3G and A3F proteins (APOBEC3 proteins) (6-8). Without Vif present, APOBEC3 proteins interact with the HIV virion and inhibit viral replication by deamination, turning cytidines to uridines in the viral DNA (9). Vif forms a complex with the APOBEC3 proteins, which serves as a substrate for binding with Cul5-E3 ubiquitin ligase to polyubiquitinate and degrade APOBEC3 proteins (10). In order to fold and form a stable structure, Vif interacts with different proteins: Elongin-B (EloB), Elongin-C (EloC) at its C-terminus, and core-binding factor subunit-β (CBFβ) at its N-terminus (11-13), forming the VCBC complex. Another protein, Cul5, interacts with Vif to form the VCBC-Cul5 complex and provides more stability and structure for the complex (13). The crystal structure of the VCBC-Cul5 complex has been solved (13). It has been shown that Vif, in the VCBC complex, has a large (α/β) domain and small (α) domain connected by a flexible linker (13,14). The linker part of Vif is important for the binding of Cul5 and for folding and creating a substrate for the E3 ubiquitin ligase complex (15).

In the first reports of the dynamics of VCBC-Cul1 and Cul5 complexes, using molecular dynamics (MD) simulations (16-18), the authors found a set of conformations that were different from the crystal structure that was previously reported (13), and showed that the Vif-ubiquitination complex is flexible. Recently (14), using MD simulations, investigators characterized and compared the dynamics of VCBC and VCBC-Cul5 complexes. They demonstrated the global dynamics of the VCBC complex but found the VCBC-Cul5 complex less flexible. The authors also revealed the efficient binding of VCBC with single-ssDNA and found out that ssDNA stimulates the formation of VCBC dimers.

Here, we applied time-lapse high-speed atomic force microscopy (HS-AFM), which has been shown to be successful for studies of the dynamics of proteins and protein-DNA complexes (19-22). Our data show that VCBC in complex with DNA is very dynamic and sampled different conformational states with the most favorable ones, including globular, dumbbell, and triangle structures. Based on the data obtained for VCBC in complex with DNA, we identified free VCBC in the same experiments, just dissociated from the VCBC-DNA complex, and followed its dynamics. We revealed that free VCBC samples conformational states and undergo transitions between globular, dumbbell, and triangle structures. The frame-by-frame analysis of time-lapse HS-AFM experiments allowed us to characterize these structures and compare dynamics for VCBC in VCBC-DNA complexes with free VCBC.

## Results

### Conformational states of VCBC in complex with DNA

As shown previously (14,23), VCBC forms a stable complex with ssDNA. This property of VCBC allows us to apply the hybrid DNA approach we used for the unambiguous characterization of the dynamics of ssDNA binding proteins using this HS-AFM methodology (22,24,25). A schematic presentation of the complex formation between VCBC and hybrid DNA is presented in Figure S1. After formation of the complex between VCBC and hybrid DNA (Figure S1 (1)) the VCBC appears on the AFM image as a dsDNA fragment with the protein sitting at the end (Figure S1 (2)). Among the four-protein complex, only Vif is capable of binding ssDNA part of hybrid DNA as it was shown in (14,23). Figure 1A presents a cartoon for VCBC bound to ssDNA where EloB, EloC and CBF-β are indicated by grey colors, and Vif, which is bound to ssDNA (bright blue line), is marked as red. The dark blue line illustrates the dsDNA part of the hybrid DNA construct.

**Figure 1.**
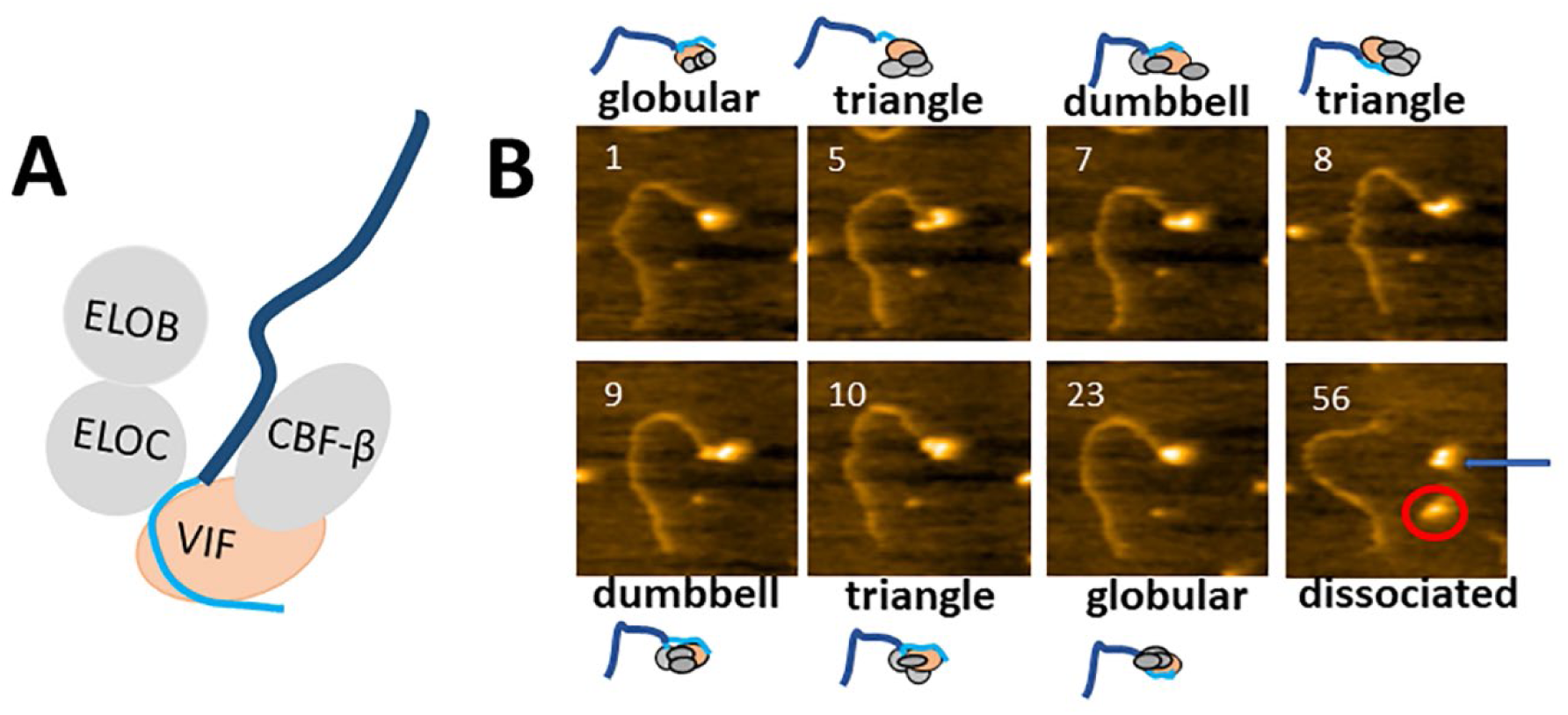
The dynamics of VCBC in the complex with DNA. **A.** Cartoon of the VCBC complex with Vif bound to ssDNA, part of hybrid DNA (bright and dark blue colors are ssDNA and dsDNA parts of hybrid DNA, respectively). **B.** Selected frames for VCBC-DNA complex from movie 1. Frame 1 presents the complex in the globular structure, which transforms into the triangle structure on frame 5, then changes its structure between dumbbell (frames 7, 9) and triangle structures (frames 8,10), forming the globular structure in frame 23, as schematically illustrated at the top and bottom of the AFM frames. Frame 56 shows dissociation of the VCBC from the VCBC-DNA complex forming a dumbbell structure (pointed by blue arrow). The protein below, marked by a red circle, is a newly assembled VCBC from nearby proteins as it is seen from movie 1. Scan size is 150 nm. Scan rate is 800 ms/frame.

To visualize the dynamics of VCBC in the VCBC-DNA complex, we applied time-lapse HS-AFM imaging. The imaging was done in the VCBC binding buffer without drying of the sample (see details of sample preparation in Experimental procedures section). After a complex of interest was found on the AFM image during scanning, we followed frame-by-frame (800 ms/frame) conformational changes in this complex, and then assembled frames into movies. From all recorded movies for VCBC in VCBC-DNA complexes, we observed several conformational states. Three conformational states of VCBC were identified as the most favorable. These states were described as follows; a compact, globular conformation, when all four VCBC proteins are close to each other; an extended, dumbbell conformation, with two clearly separated blobs; and a triangle conformation with three clear protein blobs. The transitions between all these VCBC structures are clearly seen in all collected movies (see Supporting Information).

Figure 1B presents selected frames from movie 1 (Supporting Information, Movie 1) which shows different VCBC structures and transitions between them. Specifically, frame 1 shows the globular structure of VCBC, with four proteins close to each other as schematically shown in the cartoon above the frame. The transformation of the VCBC complex from a globular structure into three visible blobs, denoted as a triangle structure, can be seen in Frame 5. Note that proteins in the triangle structure undergo some rearrangements, showing different blob sizes and distances between them, as observed when comparing frames 5, 8, and 10. Next, in frames 7-10, the complex transitions back and forth from the triangle structure to the dumbbell structure. The complex reforms the globular structure in frame 23, which remains until the complex dissociates in frame 56. Appropriate cartoons above these frames illustrate possible arrangements and rearrangements of proteins in the VCBC complex. Another interesting example of the structural transitions that VCBC undergoes is seen from selected frames in Figure 2. Again, three different structures, globular, triangle, and dumbbell are seen from this movie (Supporting Information, movie 2). Frame 11 illustrates a globular structure, which transforms into dumbbell on frame 15, then into triangle on frame 22, stays as triangle on frame 23, showing different distances between three protein blobs. Frames 23, 24, 26, and 27 show the transformation of VCBC triangle structure with three protein blobs (frame 23) into four protein blobs (frames 24, 26) and reassembly of four proteins back into the triangle structure on frame 27. The observation of such assembly and reassembly of VCBC proteins confirms, without a doubt, that globular, dumbbell, and triangle structures include all four proteins. Thus, the recorded movies confirm that the VCBC in complex with DNA is a fully assembled complex containing all four proteins.

**Figure 2.**
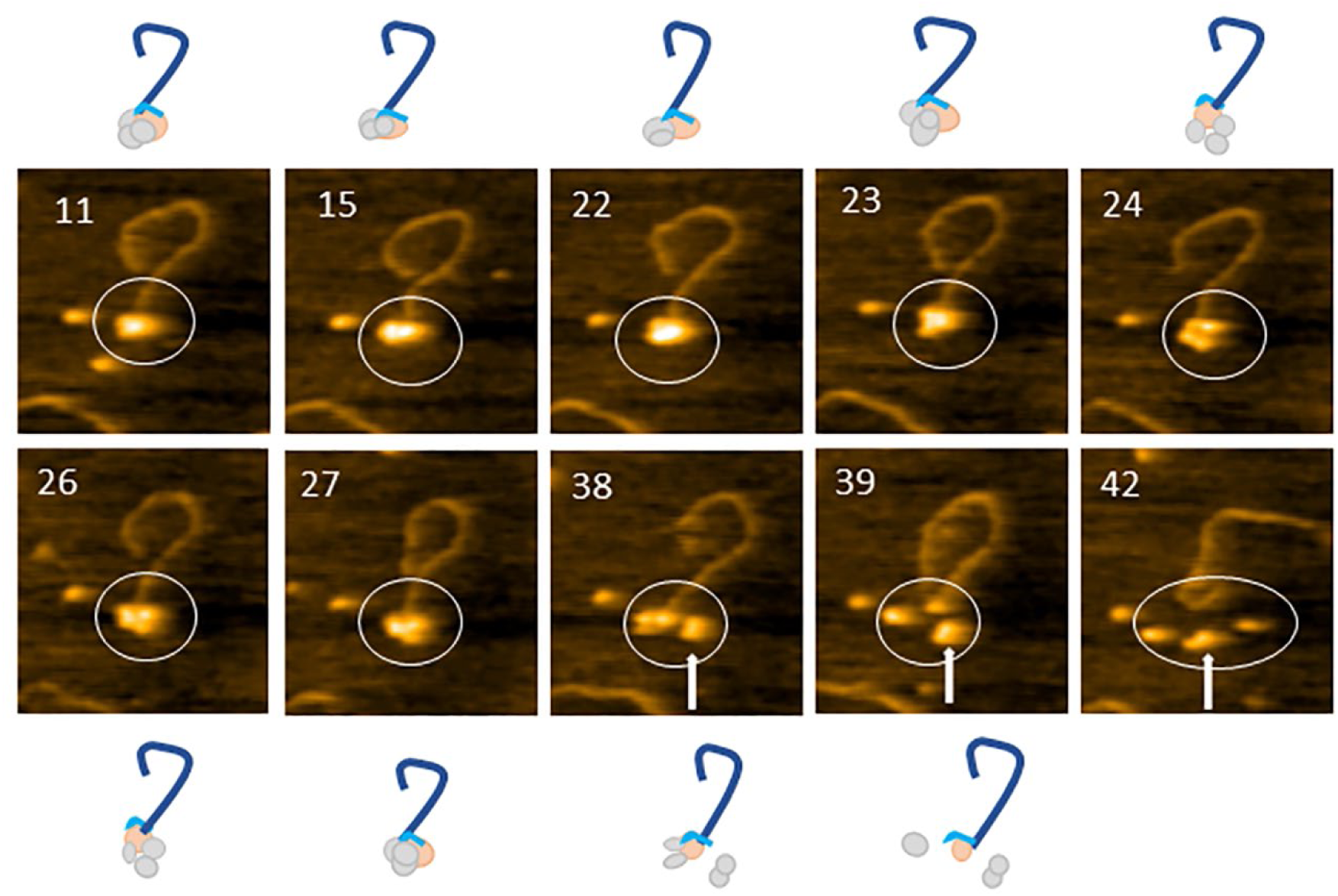
The dynamics of VCBC in the complex with DNA. Selected frames for VCBC in complex with DNA from movie 2. Frame 11 shows the globular structure of the complex, which transforms into the dumbbell structure in frame 15 and the triangle structure in frames 22 and 23. The four-protein complex can be seen in frames 24 and 26, which rearranged into the triangle structure in frame 27. Frames 38 and 39 demonstrate partial dissociation of the complex (white arrow shows two proteins together, dissociated from ssDNA). All changes in the structure of VCBC in the VCBC-DNA complex are schematically illustrated at the top and the bottom of the AFM frames. Full dissociation is shown in frame 42. Note the clearly seen part of ssDNA in frame 42 at the end of dsDNA duplex. Scan size is 150 nm. Scan rate is 800 ms/frame.

### Conformational states of free VCBC

To follow the dynamics of free VCBC, we used the AFM images for the VCBC in complex with DNA and selected a VCBC complex that was dissociated from the VCBC-DNA complex as one particle and followed its dynamics. Analysis of recorded movies shows that free VCBC is also highly dynamic and, similar to VCBC in the VCBC-DNA complex, undergoes multiple conformations, including globular, dumbbell, and triangle. Figure 3 shows one of the examples of the four-protein VCBC complex, which adopted a triangle structure as seen in a cartoon (A) and selected frames (B) from movie 3 (Supporting Information movie 3). Small cartoons above and below selected frames demonstrate the dynamics of the triangle structure of free VCBC. The rearrangements of the proteins in the VCBC complex and the changes in the distances between protein blobs in the triangle structure are seen from the presented frames. The additional selected frames from the movies are shown in the Supporting Information (Figures S2, S3). In Figure S2, the dissociation of VCBC from DNA occurs between frames 1 and 5. Free VCBC has a triangle structure on frames 5 and 14, and then adopts a dumbbell structure on frame 38 before going back to a triangle structure at the very end of the movie (Supporting Information, movie 4). Figure S3 illustrates the selected frames for free VCBC from movie 5. Frame 7 and 10 show a globular structure of VCBC, which transforms into dumbbell on frame 29, and looks like rotated dumbbell structure on frame 42.

**Figure 3.**
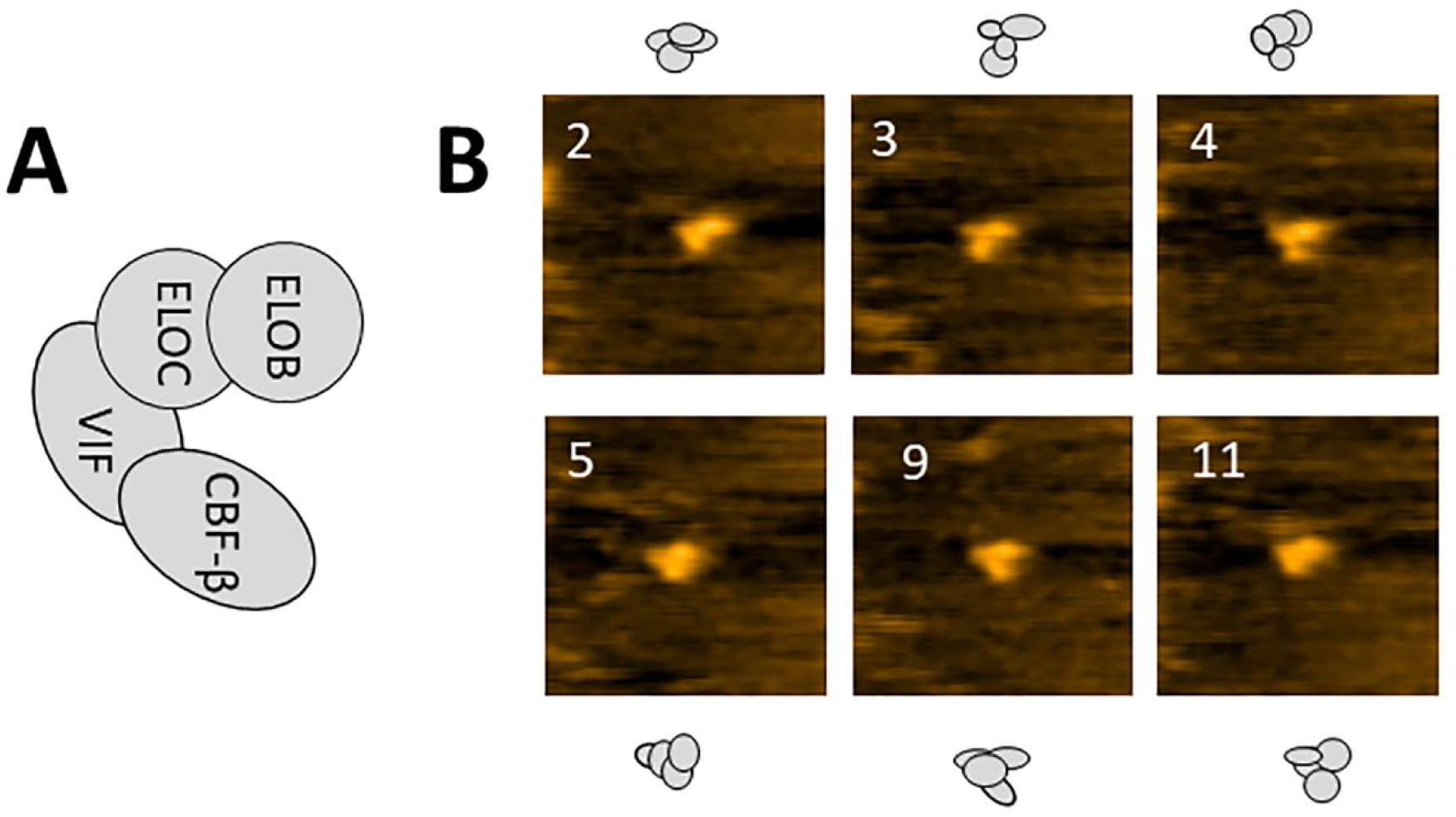
The dynamic of free VCBC. **A.** Cartoon of the free VCBC complex. **B.** Selected frames illustrating the dynamics of the triangle structure of free VCBC (movie 3) as schematically shown at the top and bottom of the AFM frames. Scan size is 150 nm. Scan rate is 800 ms/frame.

## Discussion

### Comparison of structures of VCBC in VCBC-DNA complex and free VCBC

Our time-lapse HS-AFM imaging showed highly dynamic conformational changes of free VCBC and VCBC in complex with DNA. From all recorded movies, we observed different conformational states of free VCBC and VCBC in the VCBC-DNA complex and arranged them into the three most populated structures: triangle, dumbbell, and globular structures. Frame-by-frame analysis of dozens of movies of free VCBC and VCBC in VCBC-DNA complexes allowed us to identify some characteristics of these structures.

We estimated the yield of compact, of the extended and triangle conformational states and the results are presented in Table 1. For VCBC in VCBC-DNA complexes, the yields of all three structures show an even distribution. For free VCBC, the yields of globular and dumbbell structures are similar, however, the yield of the triangle structure is higher. This indicates that for free VCBC, the triangle structure is slightly more populated when compared to the others. Comparison of the yields of all three structures between free VCBC and VCBC in complex with DNA shows similar yields for globular and dumbbell structures. However, the yield of triangle structure is slightly lower for VCBC in the complex with DNA compared to free VCBC, which indicates that DNA may affect the assembly of the triangle structure of VCBC.

**Table 1.** The yields of different structures for VCBC and VCBC in complex with-DNA.

| **Yield of structures** | **Globular** | **Dumbbell** | **Triangle** |
| --- | --- | --- | --- |
| **VCBC** | **27±9%** | **24±9%** | **49±12%** |
| **VCBC in complex with DNA** | **32±7%** | **33±8%** | **34±10%** |

In addition to the yield of the triangle structure, we followed the dynamics of VCBC in complex with DNA and compared it with the dynamics of free VCBC. The plot in Figure 4A demonstrates the changes of the angle between two arms of the triangle structure for VCBC in the VCBC-DNA complex. White lines and the arrow on AFM frame (Figure 4C) indicate the measured angle. A similar plot for free VCBC and the corresponding AFM frame is presented in Figure 4B, D. As plots in Figure 4A and B show, there is a large fluctuation between angle of the arms in the triangle structure, which may reach up to 70 degrees for both free VCBC and VCBC in the VCBC-DNA complex. Note that the triangle structure observed in our study for VCBC in the VCBC-DNA complex and free VCBC are reminiscent the U-shape structure resolved from X-ray crystallography of the VCBC-Cul5 complex (13) and the clamshell state for VCBC complex observed in MD simulations studies (14). In general, the fluctuations of the angle we observed between the arms of the triangle structure correlates with the clamshell closing and opening, as is shown in (14), and supports the significant dynamics of this structure. Another presentation demonstrating the dynamics of the triangle structure is shown in Figure S4 for VCBC in the VCBC-DNA complex (A) and free VCBC (B). Figure S4 C is the histogram of the measured angle collected from 12 movies for VCBC in the VCBC-DNA complex and fitted with a Gaussian curve. The histogram for free VCBC, collected from 15 movies, fitted with a Gaussian curve is presented in Figure S4 D. The maxima in these histograms after Gaussian fitting are similar for both free VCBC and VCBC in complex with DNA, however, the histogram for VCBC in the VCBC-DNA complex is wider than for free VCBC, which translates into a more dynamic structure for VCBC in complex with DNA. These results suggest that DNA does not restrict the dynamics of VCBC in complex with DNA. In addition to opening and closing of the triangle structure, authors in (14) provided the distances between proteins in the VCBC complex. Their measurements show the distance between EloB 80 and CBF-β 37 is around 7 nm and the distance between EloB 58 and CBF-β 37 is between 2 nm and 5 nm for the VCBC complex. HS-AFM imaging does not allow for the identification of individual proteins in the VCBC complex. However, we can measure the distances between visible blobs in the VCBC complex when they are clearly separated from each other. One of the examples of such measurements is presented in Figure S5. The distances measured for the triangle structure of VCBC in the VCBC-DNA complex (A) and free VCBC (B) are in the range of 8-10 nm, which is larger than observed from MD simulations results (14). Note that the distances between protein blobs in free VCBC and in the VCBC-DNA complex are close to each other, which emphasizes that DNA does not restrict the dynamics of the four proteins in the VCBC complex.

**Figure 4.**
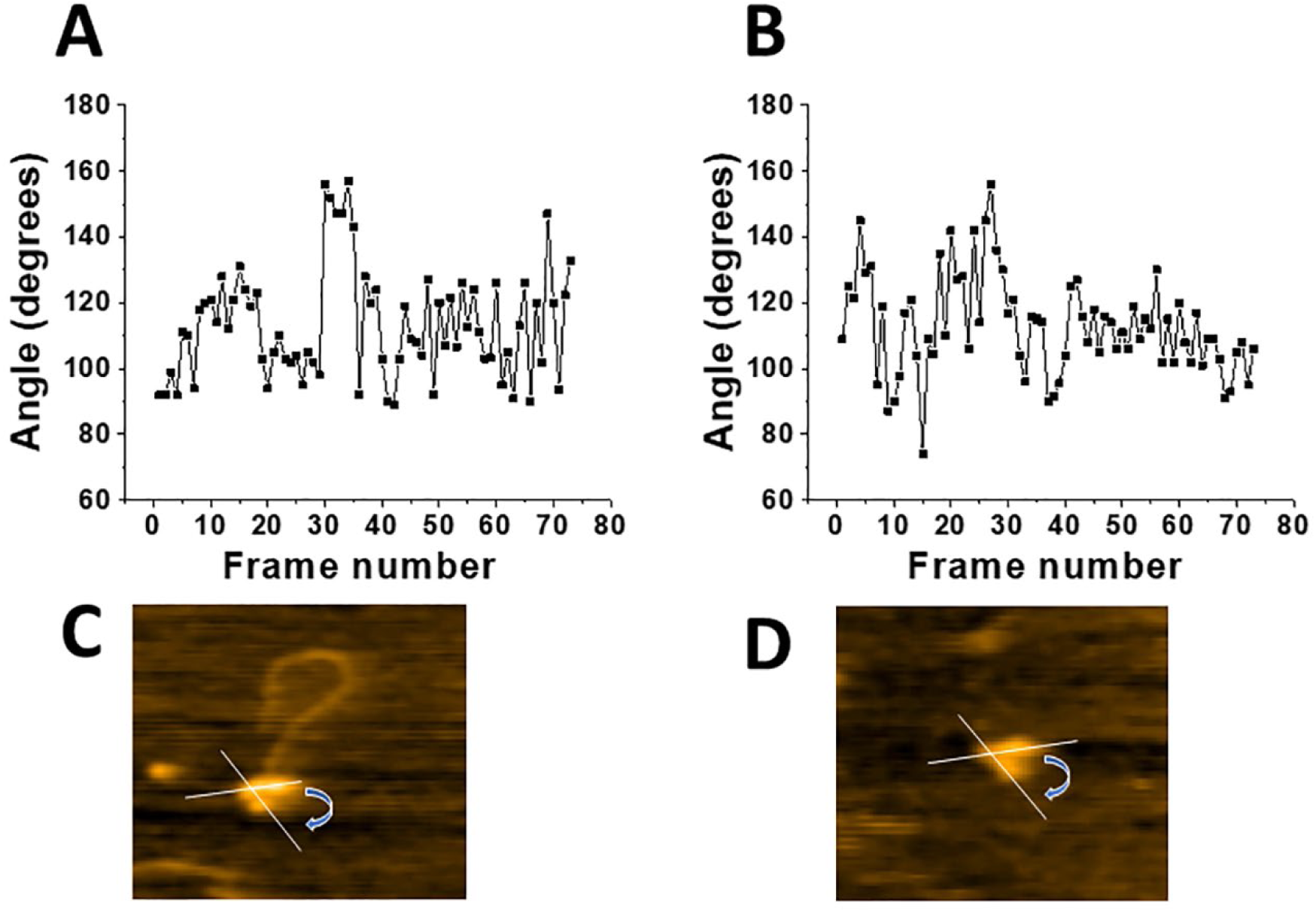
The fluctuations of the angle between arms in triangle structure. Plots show fluctuations of the angle between arms in the triangle structure for VCBC in complex with DNA (A) and free VCBC (B). AFM images of the triangle structure for VCBC in complex with DNA (C) and free VCBC (D) with the white lines and arrows illustrating the angle measurements. AFM frame sizes are 150 nm.

To follow the dynamics of the dumbbell structure, we measured the distance between the two clearly seen blobs and the results are shown in Figure 5A for VCBC in the VCBC-DNA complex and in Figure 5B for free VCBC. The blue lines on the AFM frames illustrate the measurements of the distance for VCBC in the VCBC-DNA complex (C) and free VCBC (D), respectively. Rather large dynamics for both free VCBC and VCBC in the VCBC-DNA complex can be seen for the dumbbell structure, as shown in Figures 5A and B. In this structure, two protein blobs of different sizes reflect a rather complex arrangement: the four proteins of VCBC are compacted into only two visible blobs. For this structure, the distance between two protein blobs shows fluctuation in the range of 4-14 nm, which demonstrates even larger dynamics of dumbbell structure than the triangle structure. It is an interesting observation, which may indicate that the dynamics of VCBC depend on the assembled structure. Another presentation demonstrating the dynamics of the dumbbell structure is shown in Figure S6 for VCBC in the VCBC-DNA complex (A) and free VCBC (B). Indeed, Figure S6 demonstrates AFM frames for VCBC in the VCBC-DNA complex (A), free VCBC (B), and assembled histograms (C) and (D) from collected movies. The comparison of these histograms clearly shows the difference in the dynamics of VCBC: the range of distances is wider for VCBC in complex with DNA compared to free VCBC. These data suggest that in the case of the dumbbell structure, DNA not only does not restrict, but even slightly facilitates the dynamics of VCBC in the VCBC-DNA complex.

**Figure 5.**
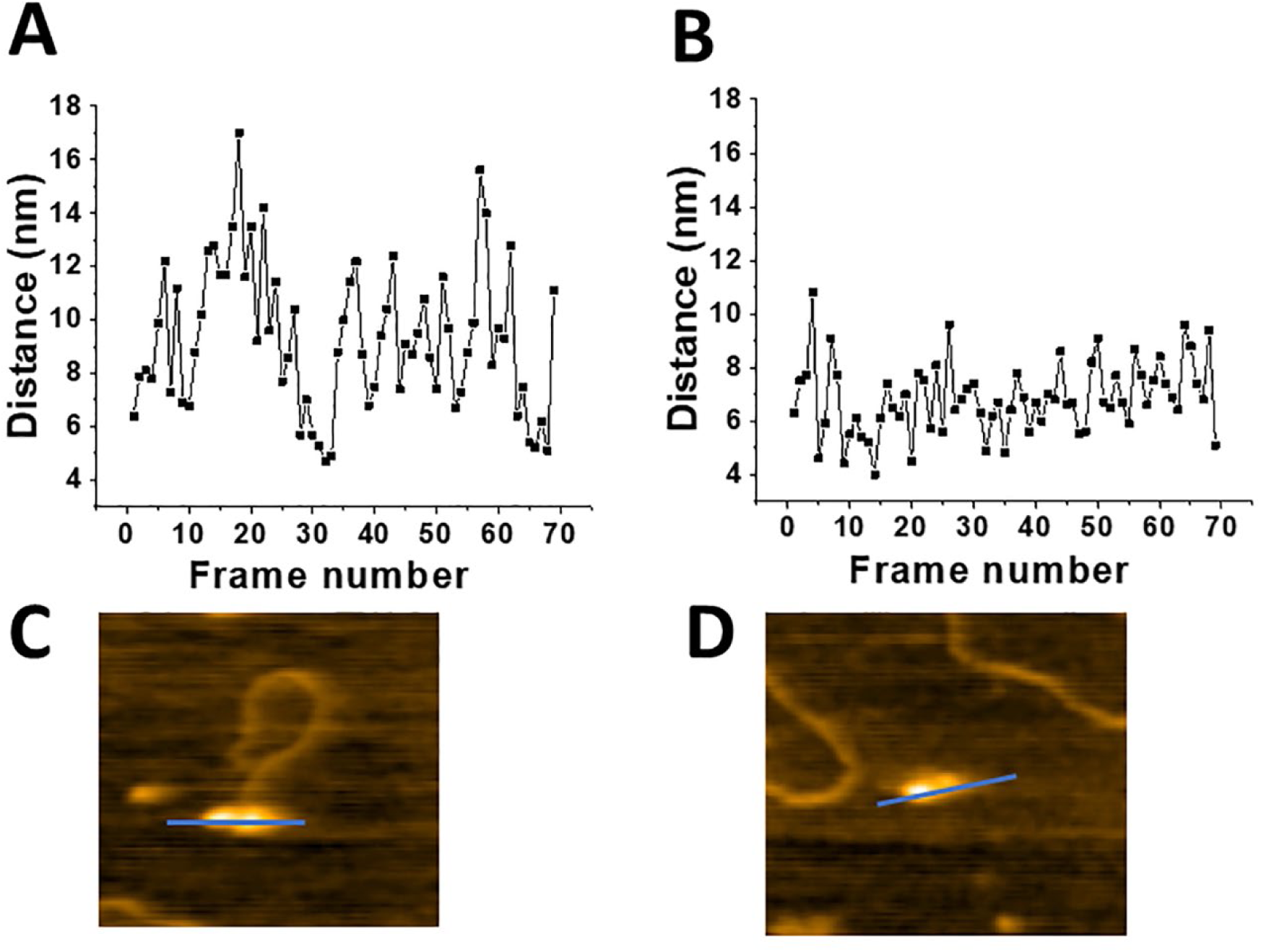
Fluctuation of the distance between protein blobs in the dumbbell structure. Graphs show the fluctuation of the distance between two blobs in the dumbbell structures for VCBC in complex with DNA (A) and free VCBC (B). B and D are AFM images of the dumbbell structure for VCBC in complex with DNA and free VCBC. Blue lines and arrows show the method of measurement. AFM frames sizes are 150 nm.

We also followed the dynamics of the globular VCBC structure for both free VCBC and VCBC in the VCBC-DNA complex. The results of the estimated volumes for this structure are presented in Figure S7. The data are assembled into histograms and the standard deviations, calculated after fitting with Gaussian curves, demonstrated the dynamic behavior of this structure. However, the most interesting fact is that VCBC in the VCBC-DNA complex has two separate peaks in the histogram (Figure S7 A) compared to only one peak for free VCBC (Figure S7 B). The value of the maximum for the volume in the case of free VCBC (318 ± 80 nm^3^) is close to the first maximum in the histogram for VCBC in the VCBC-ssDNA complex (298 ± 100 nm^3^). Based on this fact, we assigned the first peak in Figure S7 A and the peak in Figure S7 B to the monomeric state of VCBC. The second peak in histogram (Figure S7 A) is roughly double the value of the first peak (703 ± 80 nm^3^), which points to the possible formation of the VCBC dimers in the presence of DNA. The formation of VCBC dimers in the presence of DNA correlates with the results reported previously (14), where authors observed the formation of VCBC dimers in the presence of 14 oligonucleotides (dT14). According to the standard deviation values, the dynamics of VCBC monomers in the VCBC-DNA complex and free VCBC are close to each other, suggesting that DNA does not affect the dynamics of the globular structure.

Together, the comparison of the results between free VCBC and VCBC in the VCBC-DNA complex demonstrates that DNA does not restrict the dynamics of the triangle, dumbbell, and globular structures. In general, the dynamics of VCBC in the presence of DNA is similar or slightly larger than the dynamics of free VCBC. Based on crystallographic and MD simulation data, Vif adopts a folded conformation, interacting with EloB, EloC, and CBF-β in the four-protein VCBC complex. The flexible linker located between the α/β and α domain of Vif is responsible for conformational changes of VCBC (14). Based on the fact that DNA does not affect the dynamics of VCBC and the type of VCBC conformations, we suggest that DNA binds a region away from the flexible linker of Vif.

In summary, our studies directly demonstrate for the first time the very large conformational dynamics of the VCBC complex. Among numerous intermediate conformational changes of the VCBC complex, we revealed the most populated structures, which we identified as triangle, dumbbell, and globular structures. Although our results show large conformational dynamics in all three favorable structures, we find that the most dynamic structures are the triangle and dumbbell structures, which suggests that the range of dynamics may depend on the conformational state of the VCBC complex. Also, our data not only confirmed the global dynamics of VCBC, characterized from MD simulations, but demonstrated even larger conformational mobility of VCBC, whether for free VCBC or in the VCBC-DNA complex. Our results will provide a better understanding of the structure and dynamics of the VCBC complex and may accelerate the progress in this field.

### Experimental procedures

#### VCBC protein

The Hxb2 VCBC used in this study was obtained from Dr. J. Gross (UCSF) and purified as described in (26).

#### Hybrid DNA substrate

Hybrid DNA was assembled as described in (27) and consists of a 69nt ssDNA tail with 379 bp dsDNA as a tag (see cartoon in Fig.S1 (1)). Briefly, 89 nt long synthetic oligos were annealed with short 23 nt oligos (Integrated DNA Technology; IA) to create a short duplex with sticky ends. This short duplex was ligated with a 356 bp dsDNA fragment that was complementary to the duplex sticky ends. The hybrid DNA was gel purified (Qiagen kit) and resuspended in 10mM Tris, pH 7.5, and 1mM EDTA.

#### The preparation of VCBC-DNA sample

Hybrid DNA (60 nM) was mixed with Hxb2 VCBC (240 nM) at ratio of 1:4 in binding buffer, containing 50 mM HEPES, pH 7.5, 100 mM NaCl, 5 mM MgCl_2_, and 1mM DTT. After the incubation of the mixture at 25 °C for 15 min, the sample was diluted up to 2 nM with the binding buffer and immediately deposited on a functionalized mica surface (28,29).

#### HS-AFM imaging

The detailed description of time-lapse HS-AFM imaging is described in (21,22,30). In brief, two microliters of diluted VCBC-DNA complex were deposited for 2 min on a mica surface and washed with binding buffer. The HS-AFM scanning was initiated immediately after thee washing step without drying the sample. The scan size was 300 nm x 300 nm. After the complex of interest was selected, frame-by-frame recording was started with a scan rate of 800 ms/frame. The probes for imaging were grown under an electron beam using short cantilevers (BL-AC10DS-A2, Olympus, Tokyo, Japan) with spring constant 0.1-0.2 N/m^-1^ and resonance frequency of 400-1000kHz.

#### Analysis of the HS-AFM data

Movies were assembled after frame-by frame recording of VCBC in the VCBC-DNA complex and free VCBC. Analysis of the observed structures was performed by measuring the yield of different structures and their structural characteristics. For globular structures, the volume of the VCBC in the VCBC-DNA complex and free VCBC were measured as described in (31), using the cross section feature from Femtoscan online software (Advance Technologies Center, Moscow, RU). For dumbbell structures, the same cross section feature was used to measure the distance between two clear blobs in the structure (21). For triangle structures the feature for angle measurement was used from Femtoscan software. On average, for each structure ∼10 movies were analyzed, and the data were assembled into histograms and fitted with Gaussian curves.

## Acknowledgments

We thank to Drs. John Gross and Xi Lui (USCF) for providing the VCBC protein.

## Funding and additional information

This work was supported by the NIH grant R01 GM108006 to Y.L Lyubchenko.

